# Bacterial-driven development is mediated by Calcium-Dependent Intrinsic Apoptosis in the Squid-Vibrio Symbiosis

**DOI:** 10.64898/2026.07.15.738745

**Authors:** Madison Emery, Alice Breaux Walker, Bethany Elizabeth Rader, Elizabeth Anne Chapman Heath-Heckman

## Abstract

The presence of beneficial microbes serves as a post-embryonic developmental cue in a wide array of metazoan species. However, the mechanisms through which mutualistic bacteria induce developmental processes such as apoptosis are poorly understood. A leading model system utilized to study bacteria-induced developmental apoptosis is the Hawaiian bobtail squid, *Euprymna scolopes*, whose bacterial symbiont *Vibrio fischeri* induces several developmental events upon colonization of the squid’s light organ. Upon hatching the light organ possesses ciliated epithelial fields (CEFs) and appendages that facilitate the collection of *V. fischeri* from the ambient seawater for symbiont colonization of the internal crypt spaces. To better understand the molecular pathways underpinning bacterial-induced development occurring in these appendages, we isolated appendages from hatchling (0-1h) and aposymbiotic and symbiotic *E. scolopes* light organs (18h) for RNA sequencing. Our analysis of this transcriptomic dataset indicated that symbiotic appendages undergo intrinsic apoptosis in response to excessive cytosolic calcium. Further experiments found that symbiotic appendages exhibited increased cytosolic calcium and mitochondrial membrane potential overload relative to their aposymbiotic counterparts. In comparing our appendage specific gene expression to previously published whole light organ transcriptomic dataset, we identified increased presence of apoptosis inducing factor (AIF) and decreased expression of transcripts related to protein folding in the appendages as potential mechanisms of apoptotic signal specificity to the CEF. Together, these data suggest a central role of cytosolic calcium in the developmental apoptotic signaling induced by *V. fischeri* colonization of the *E. scolopes* light organ.

**Importance:** Bacterial cues are known to induce post-embryonic animal development, but the mechanisms mediating crosstalk between bacterial product recognition and developmental dynamics require further study. Because of its binary nature and clear developmental phenotypes, the *Euprymna scolopes*- *Vibrio fischeri* system is an excellent model for symbiont-induced development. Here, we characterize the symbiont-induced apoptotic signaling that mediates the loss of *V. fischeri* recruitment structures in the *E. scolopes* light organ following colonization. Transcriptomic changes in colonized light organs and subsequent microscopy-based experiments support that acquisition of *V. fischeri* induces loss of the ciliated symbiont recruitment structures in juvenile light organs via intrinsic apoptotic signaling initiated by excess cytosolic calcium. This work advances our understanding of how mutualistic bacterial cues initiate signal transduction to induce developmental programming and may serve as a foundation upon which we can begin to disentangle the ways in which more diverse and complex microbial communities influence post-embryonic development.

## Introduction

Exposure to bacteria throughout 400 million years of animal evolution has resulted in nearly ubiquitous microbial symbioses across the Metazoa (1, 2). Because of the intimacy of these interactions, symbiotic bacteria often have profound impacts on the biology of their animal hosts. One impact that is commonly seen across diverse animal phyla is host development being shaped by their microbial symbionts (2–4). Symbiotic bacteria have been shown to induce larval settlement and morphogenesis in marine invertebrates (4–7), as well as shape organ and organ system development (2, 3, 8, 9). While external environmental microbes most commonly induce settlement and morphogenesis, organ and organ system development are typically shaped by internal symbionts within a consortia making specific mechanisms of bacteria-stimulated development difficult to identify (2–8, 10, 11). As such, binary symbiotic systems can serve as powerful tools for understanding how bacterial symbionts shape organ development (11, 12).

One of the leading model systems utilized to study bacteria-induced post-embryonic development is the binary relationship between the Hawaiian bobtail squid, *Euprymna scolopes*, and its bioluminescent bacterial symbiont, *Vibrio fischeri* (*12, 13*). *V. fischeri* reside extracellularly within the internal crypt spaces of the *E. scolopes* specialized symbiotic organ, known as the light organ (Figure 1A-B). At the time of hatching, the light organ has external ciliated epithelial fields (CEFs), which form paired, symmetric epithelial appendages (Figure 1B) (9). The light organ cilia facilitate symbiont acquisition from the surrounding seawater via the generation of flow fields and secretion of mucus containing molecules such as nitric oxide (NO) that select against non-*V. fischeri* bacteria (14, 15). The process of symbiont acquisition occurs in the first three to four hours after hatching and occurs only once within the squid’s lifespan. As such, after *V. fischeri* successfully colonizes the light organ, the appendages and the CEFs used to facilitate colonization undergo apoptosis (9, 16).

**Figure 1.**
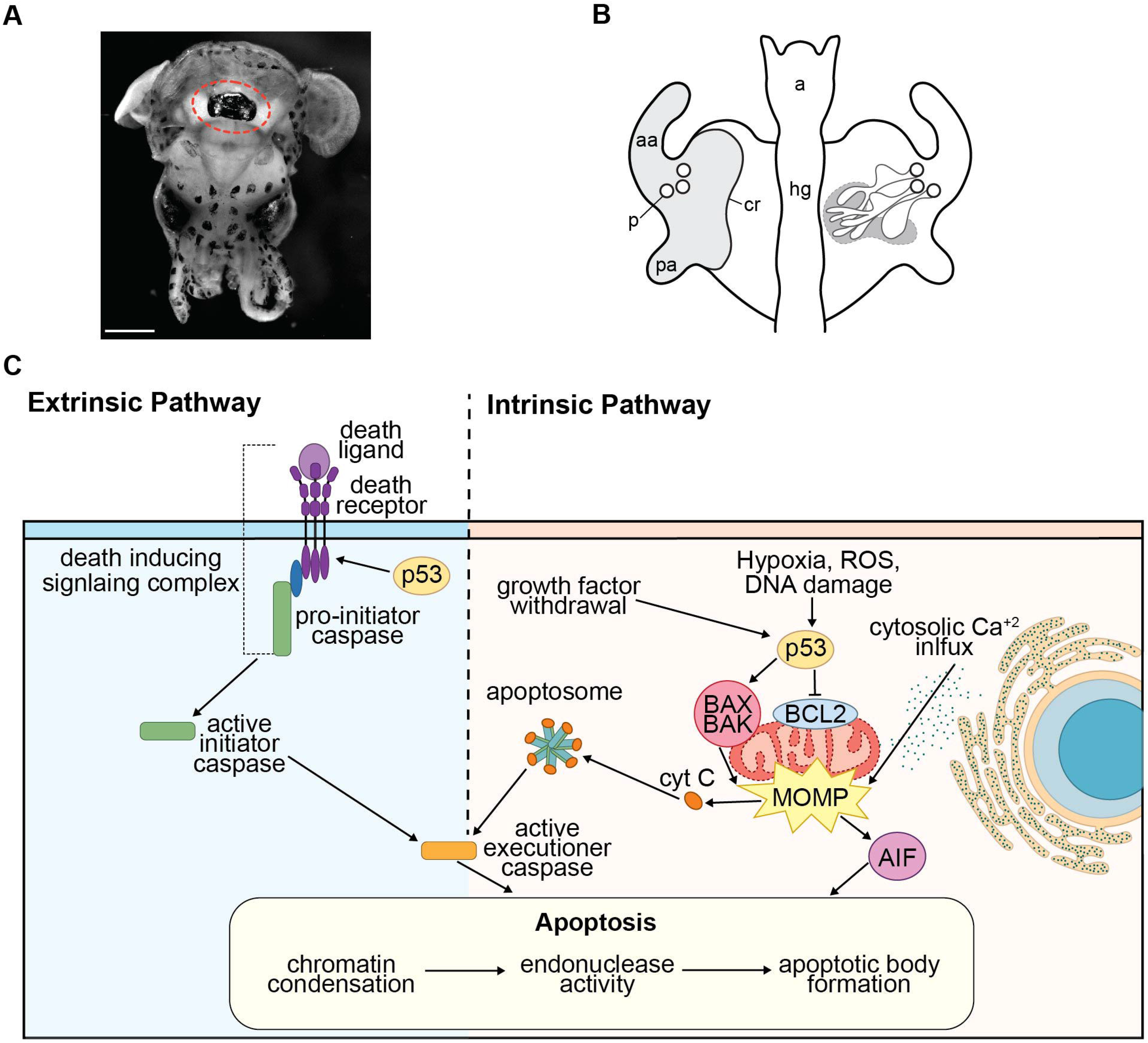
Overview of the *E. scolopes* light organ and apoptotic pathways (A) Whole-mount dissection of hatchling *E. scolopes* with the light organ and surrounding ink sac circled (B) diagram of hatchling *E. scolopes* light organ adapted from Walker et al. 2026 (87). External structures are depicted on the left side of the diagram, with gray shading indicating the ciliated epithelial field (CEF). Internal structures are depicted on the right side of the diagram. Deep crypt spaces where *V. fischeri* are housed are encircled. Abbreviations: aa: anterior appendage, pa: posterior appendage, p: pore, cr: ciliated ridge, a: anus, hg: hind gut. (C) Diagram of extrinsic and intrinsic apoptotic pathway components present in the *E. scolopes* genome and previously studied apoptotic events in CEF loss/appendage regression in chronological order. Adapted from Vroom et al. 2022.

While bacterial-induced development in this system is well documented, only some parts of the signaling cascade regulating this process are known. Loss of the appendages and CEFs is initiated by host recognition of both lipopolysaccharide and peptidoglycan, two components of the gram-negative cell envelope. Exposure to the combination of these two bacterial microbe-associated molecular patterns induces similar levels of apoptosis to what is observed in the CEFs of light organs colonized by *V. fischeri* (17). However, much remains unknown about what pathways are involved in transducing the apoptotic signal to the appendages/CEF following recognition in the internal crypt spaces (18, 19). Within the CEF chromatin condensation is decoupled from the subsequent stage of apoptosis, DNA fragmentation via endonucleases (16, 18, 19). It has been shown that NO levels must attenuate for chromatin condensation, an early cellular event that occurs during apoptosis, to occur (19). While chromatin condensation is reversible by removing *V. fischeri* from the light organ with antibiotics up until 12h post hatching, endonuclease activity occurs after this timepoint and is not reversible nor is it directly tied to NO attenuation (16, 18, 19).

It is unknown what signaling pathways are involved with the initiation of later stage apoptosis following NO attenuation. There is some evidence supporting the involvement of p53 family transcription factors (20). However, this involvement does little to narrow down what apoptotic singling pathway is responsible for loss of the CEF in the *E. scolopes* light organ.

There is genomic evidence that *E. scolopes* are capable of intrinsic and extrinsic apoptotic signaling, both of which can be mediated by p53 transcription factors (21, 22). Extrinsic apoptosis is mediated by transmembrane death receptors, which upon binding their respective pro-death ligands recruit adaptor proteins and pro-initiator caspases to form signaling complexes that are responsible for ultimately activating executioner caspases (23–25). Intrinsic apoptosis is initiated by intracellular signals, including hypoxia, excess cytosolic calcium, growth factor withdrawal, and oxidative stress. In response to these stimuli, pro-apoptotic Bcl-2 family proteins mediate the opening of mitochondrial permeability transition pore causing the loss of mitochondrial membrane potential and releasing pro-apoptotic proteins into the cytosol where they initiate either caspase dependent or caspase independent cell death (Figure 1C) (24–28).

Determining what apoptotic signaling is occurring in the CEF and appendages during symbiont induced morphogenesis in *E. scolopes* would give clues as to the steps connecting bacterial product presentation and apoptosis initiation. Therefore, we isolated appendages from freshly hatched squid along with appendages from the light organs of aposymbiotic and symbiotic squid at 18h for RNA sequencing to identify the genes and pathways differentially expressed as endonuclease activity increases in the CEF (16). Through this transcriptomic dataset and subsequent experimental validations, we provide insight into the apoptotic pathways that mediate appendage regression and mechanisms of apoptotic signal specificity in *E. scolopes* symbiont-induced morphogenesis.

## Results

On average, 35.9 million reads per replicate mapped to the filtered *E. scolopes* reference transcriptome, resulting in an average mapping rate of 77.9% (Supplemental File 1). In total, 26,161 transcripts passed the expression threshold and were used in downstream analysis. A principal component analysis (PCA) showed no obvious outliers, so downstream differential expression analysis proceeded with three replicates in each the following treatment groups: 0-3h hatchling (hatch) appendages, 18h aposymbiotic (apo) appendages, and 18h symbiotic (sym) appendages (Supplemental Figure 1).

When comparing expression of all three treatments at once, we identified 4343 significantly differentially expressed genes (DEGs). Interestingly, the expression patterns of these DEGs in the hatch replicates do not consistently mirror their expression in the apo replicates, despite the lack of *V. fischeri* exposure in both of treatments. While some DEGs show similar relative expression in hatch and in apo appendages, the majority are either an intermediate expression level between what is seen in apo and sym replicates or mirror the expression levels observed in sym appendages (Figure 2). This general trend is further supported by the number of DEGs identified in each pairwise comparison, where the most DEGs were present between sym and apo (4343), followed by apo and hatch (3859), and finally sym and hatch (2844) (Supplemental File 2).

**Figure 2.**
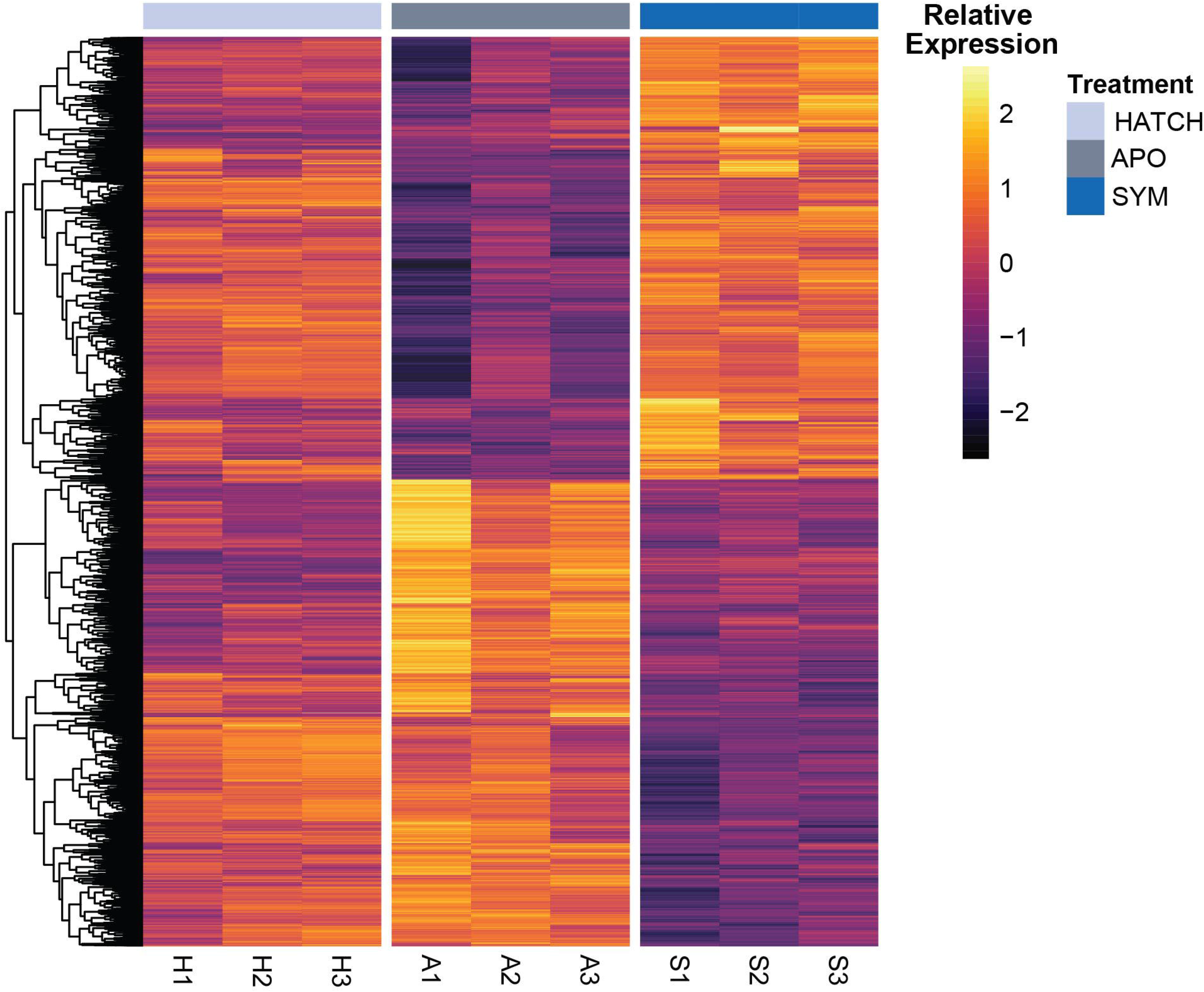
There are substantial changes in gene expression between symbiotic, aposymbiotic, and hatchling appendages. Heatmap of relative rlog expression of all DEGs between hatchling (hatch), aposymbiotic (apo), and symbiotic (sym) appendages clustered by expression similarity. Labels at the bottom of the columns indicate individual replicates.

To identify the biological processes with significant differential expression in each pairwise comparison we used GO-MWU (29). This continuous approach to identify gene ontology (GO) enrichments resulted in a large number of up and down regulated terms in each comparison (Supplemental File 3). Therefore, these results were summarized by their parent GO terms. Proportionally, the parent terms of the GO enrichments observed being upregulated in sym vs apo and in sym vs hatch were similar, with substantial upregulation of terms related to development, immunity, metabolism, the nervous system, and ion transport. However, the number of upregulated terms related to development, metabolism, the nervous system, and ion transport were two or more times higher in the sym vs apo comparison relative to the sym vs hatch comparison. Within the apo vs hatch comparison, we found evidence suggesting that perhaps the delay in development due to the lack of exposure to *V. fischeri* is a stressful condition in apo appendages, as more stress response enrichments were upregulated in apo replicates relative to hatch. The upregulation of processes related to stress responses in apo appendages is also seen in the sym vs apo comparison. However, the majority of stress response terms in this comparison are being upregulated in sym appendages relative to apo (Figure 3).

**Figure 3.**
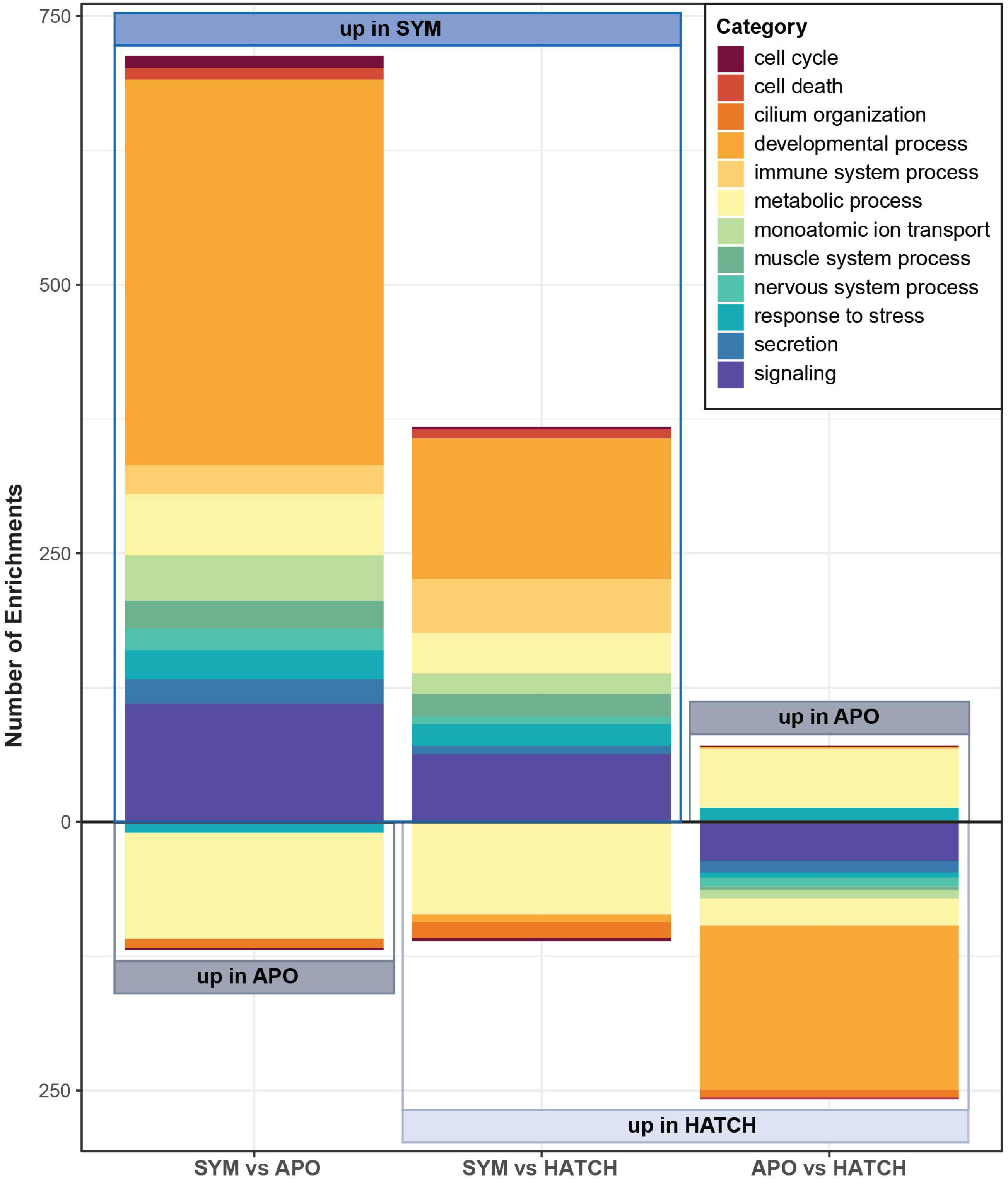
Symbiotic appendages upregulate similar categories of biological processes relative to both apo and hatch appendages, but at different magnitudes. Rank-based GO enrichments in each pairwise comparison summarized by number of enrichments with a given parent GO term.

To best leverage this appendage-specific dataset to isolate potential apoptotic pathways involved in appendage regression and discern potential mechanisms of apoptotic signal specificity to the CEFs and appendages, we reanalyzed previously published RNAseq data from Moriano-Gutierrez *et al.* of apo and sym whole light organs at 24h using the same pipeline as our appendage data (30). This comparison allows for the identification of processes regulated differently with symbiosis in the appendages relative to the rest of the light organ and has the potential to isolate a secreted signal present in both datasets that may be involved in apoptotic signal transduction. From the whole light organ RNAseq data, an average of 19.1 million reads per replicate mapped to the filtered *E. scolopes* reference transcriptome, resulting in an average mapping rate of 76.8% (Supplemental File 1). As the whole light organ dataset had lower sequencing depth than our appendage RNAseq data, it had less power to detect DEGs (31). This resulted in the identification of 857 DEGs between sym and apo whole light organs. Despite differences in sequencing depth, 336 overlapping DEGS were identified between the sym vs apo appendage DEGs and the whole light organ analysis (Supplemental file 2). Similarly, there were 95 differentially expressed BP GO terms, 39 of which were shared between enrichments identified in the comparison between sym and apo appendages. Notably, only one of these differentially regulated terms, protein folding, had opposing directionality between the two datasets. This term was upregulated in sym light organs relative to apo but down regulated in sym appendages relative to apo appendages (Supplemental file 3). Additionally, none of the differentially regulated BPs had nervous system process or ion transport parent terms, suggesting that these enrichments may be unique to the sym appendages (Figure 4).

**Figure 4.**
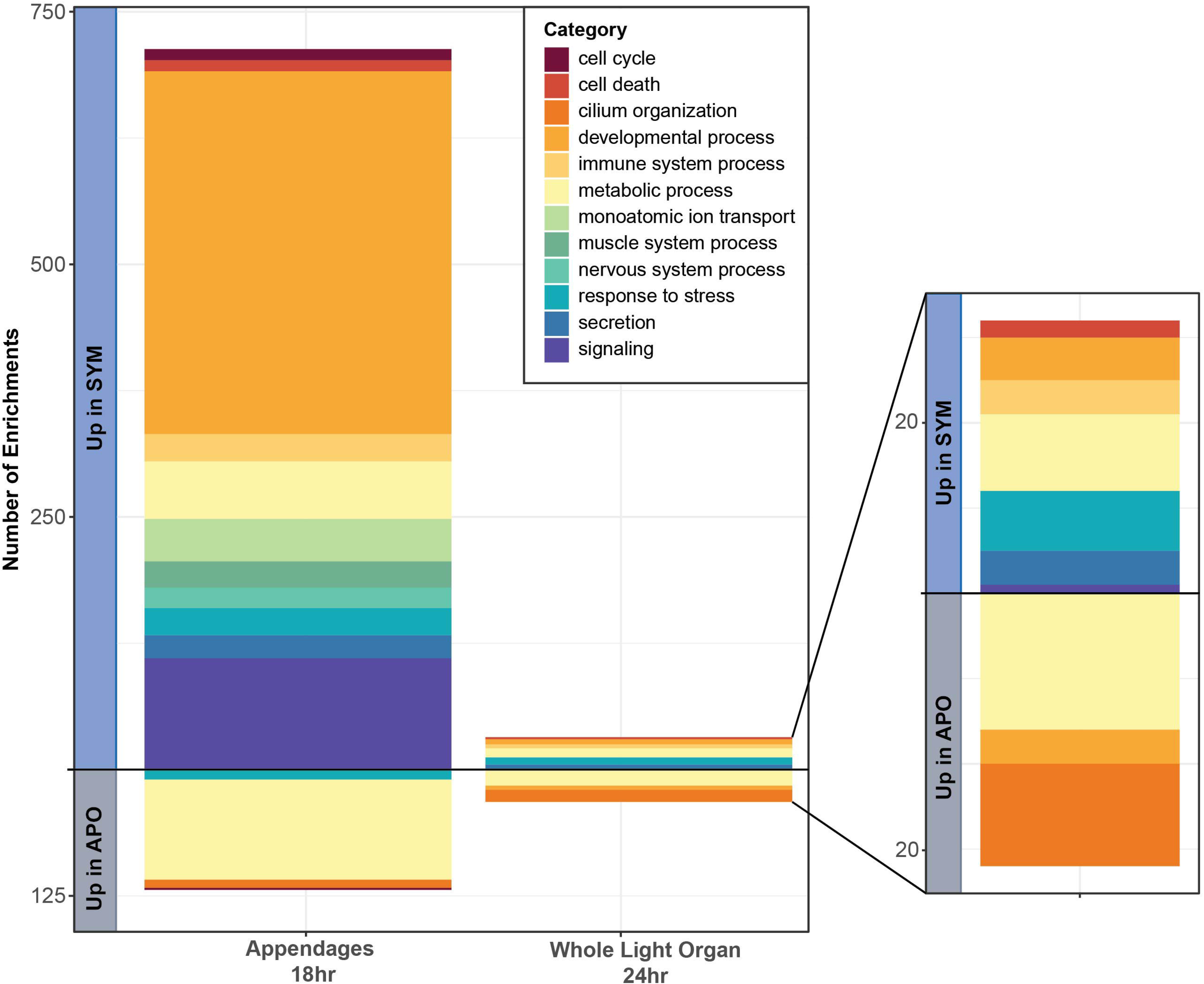
Enrichments in processes related to the nervous system and ion transport are unique to the 18h appendage dataset and are not found in whole light organs at 24h. Comparison of the number of enriched GO terms with a curated set of parent GO terms in apo vs sym replicates found in our appendage specific dataset as opposed to our reanalysis of whole light organ data generated by Moriano-Gutierrez *et al*.

The increased depth of the appendage dataset along with the inclusion of the hatch treatment group resulted in more specific insight into the apoptotic signaling present in sym appendages. The only specific apoptotic pathway GO term upregulated in sym appendages relative to both apo appendages and hatch appendages was “intrinsic apoptotic signaling pathway”. An additional upregulated term in sym appendages vs apo appendages indicates that the initiation of intrinsic apoptosis may be related to the release of calcium (Ca^2+^) from the endoplasmic reticulum (ER) (Figure 5a). There are several upregulated terms in sym appendages vs apo and hatch appendages that support this hypothesis, as they indicate changes to the concentration of cytosolic Ca^2+^ in sym appendages (Figure 5b). However, there are also terms that suggest sequestration of Ca^2+^ in the ER. Specific DEGs also follow this contradictory pattern, as two ryanodine receptors (RyRs), which mediate release of Ca^2+^ from the ER, are upregulated, along with several sarcoendoplasmic reticulum calcium ATPase (SERCA) pumps, which sequester Ca^2+^ in the ER (Figure 6a) (32, 33). Though the gene expression data are unclear as to what is occurring to cytosolic Ca^2+^ concentrations in sym appendages, there is further support for intrinsic apoptosis mediating appendage regression via the upregulation of apoptosis-inducing factor 1, mitochondrial (*aifm1*), which is a pro-apoptotic factor that is secondarily released from the mitochondrial membrane during late-stage intrinsic apoptosis. Notably, RyRs, SERCA pumps, and *aifm1* were not differentially expressed between apo and sym whole light organs at 24h (Figure 6b).

**Figure 5.**
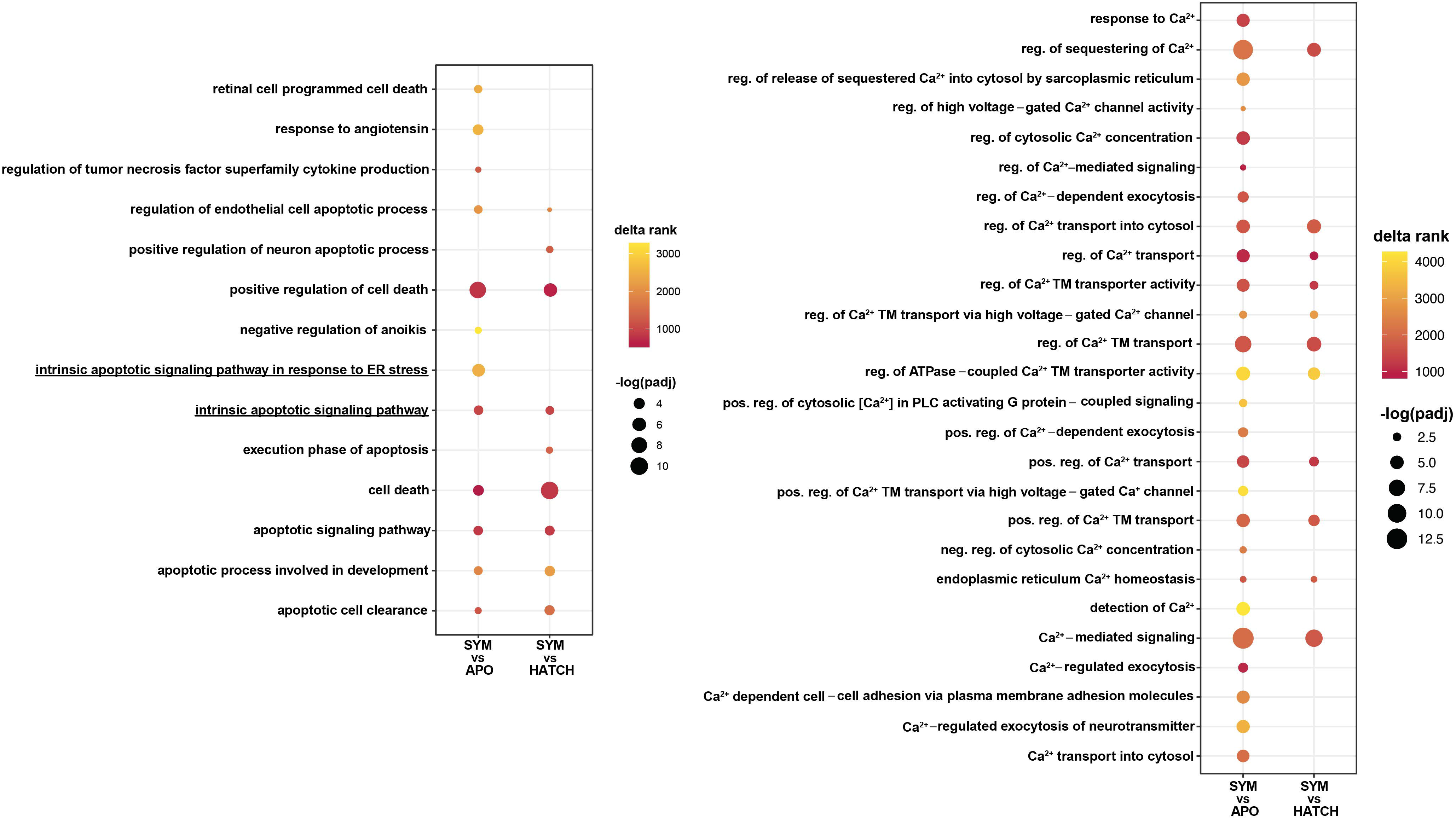
Enriched GO terms suggest that cytosolic Ca^2+^ may be inducing intrinsic apoptosis in the ciliated epithelial cells of sym appendages. Rank-based gene ontology enrichments that are upregulated in sym appendages relative to apo and/or hatch appendages related to (A) apoptosis and (B) Ca^2+^. Color of each point indicates delta rank, dot size indicates negative log-adjusted p-value. Abbreviations: reg: regulation, TM: transmembrane, pos.: positive, PLC: phospholipase C, neg.: negative.

**Figure 6.**
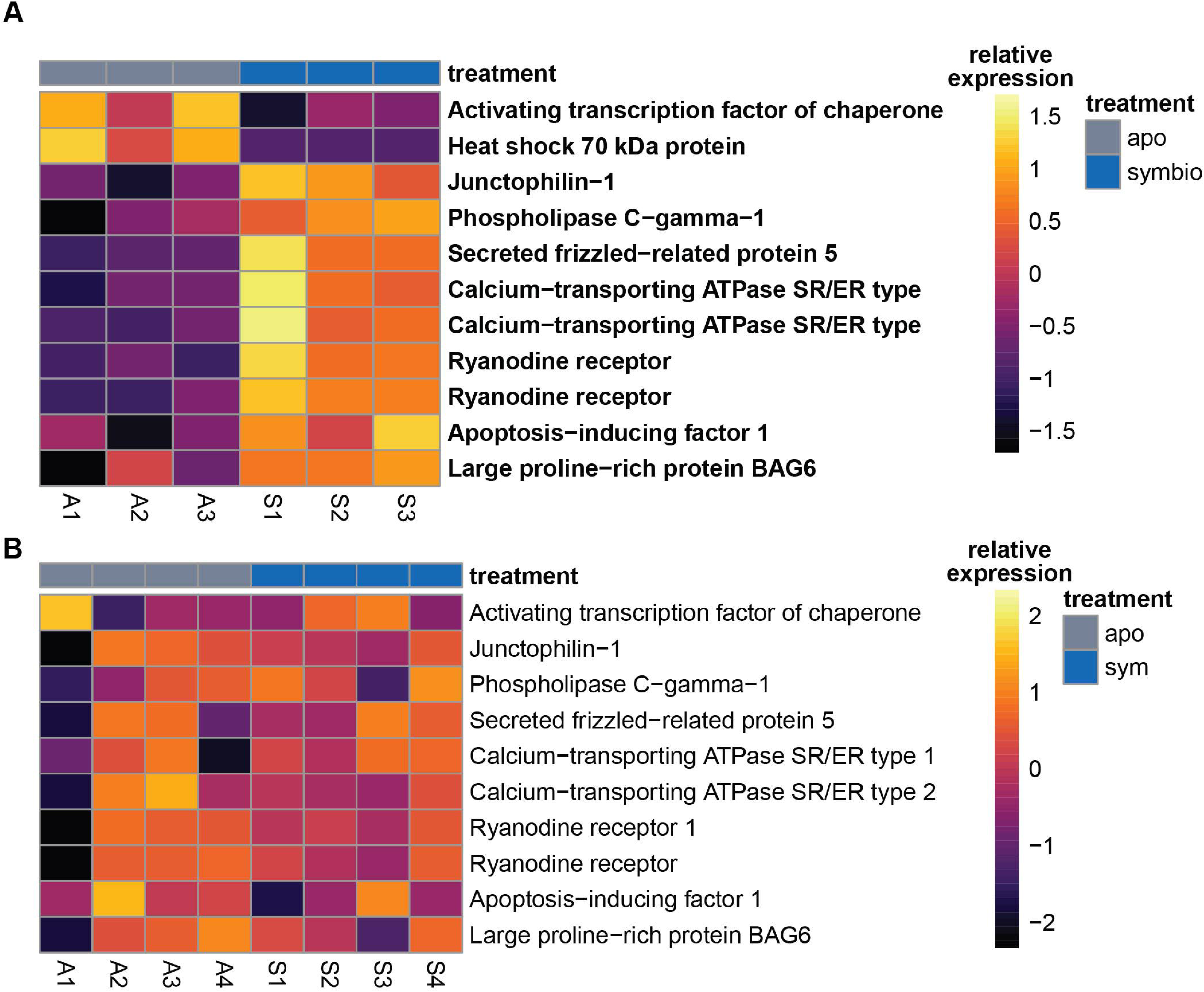
Key transcripts related to Ca^2+^-mediated intrinsic apoptosis are significantly differentially expressed in sym appendages relative to apo but are not differentially expressed in whole light organs. Heatmaps of relative expression of key transcripts supporting presence of Ca^2+^-mediated intrinsic apoptosis in (A) sym and apo appendages at 18h and (B) sym and apo whole light organs at 24h. Bolded transcript names indicate that it is a DEG. Labels at the bottom of the columns indicate individual replicates.

To test if sym appendages had higher levels of cytosolic Ca^2+^ relative to apo, we used Rhod-3, a live stain that preferentially binds cytosolic Ca^2+^. While sym appendages intermittently had distinct puncta with large amounts of cytosolic Ca^2+^ at 18h, there was no significant difference the puncta number in 18h sym appendages relative to apo. However, at 24h, sym appendages had significantly more puncta than apo (two-way ANOVA, p= 0.0002) (Figure 7).

**Figure 7.**
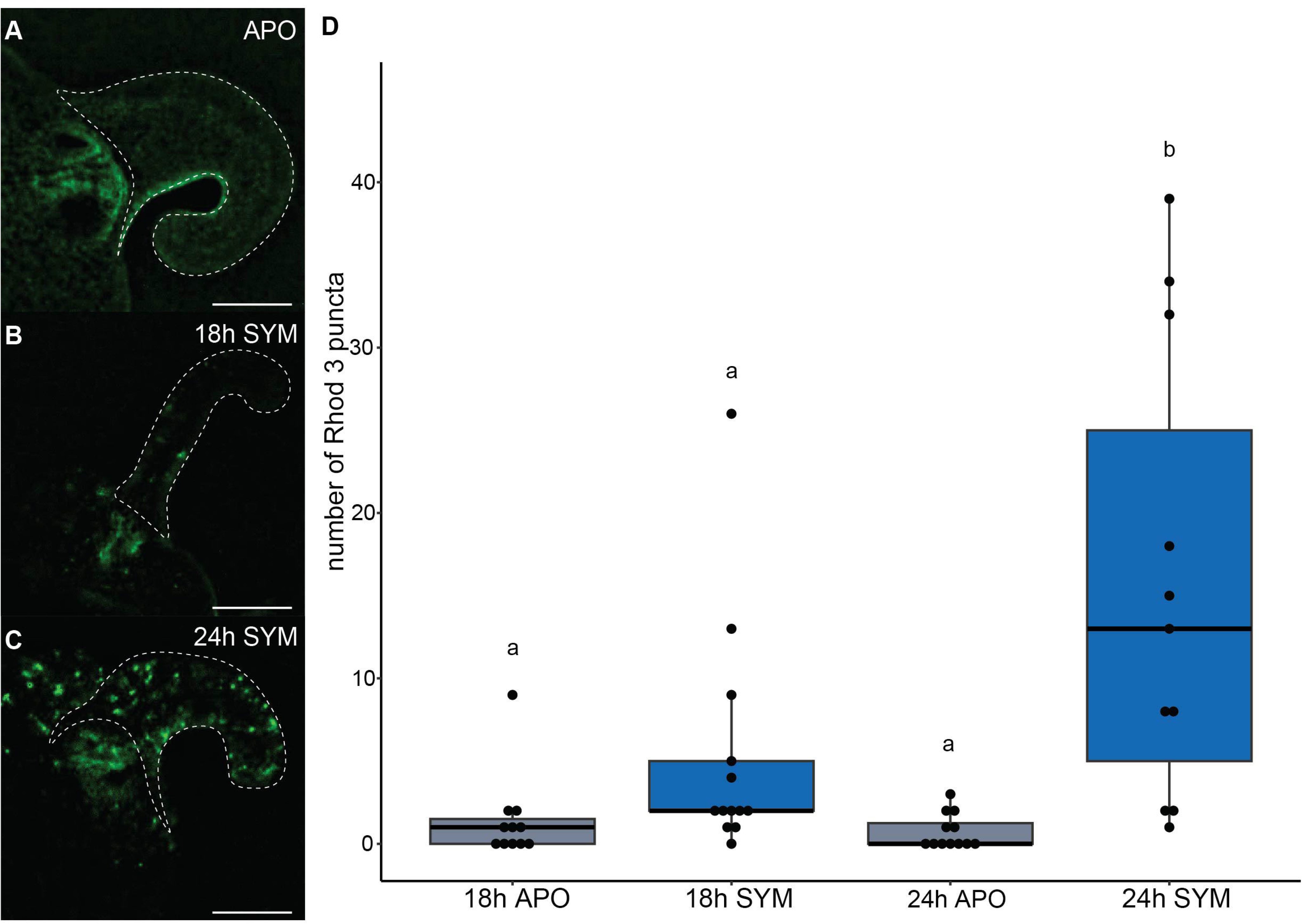
Colonization by *V. fischeri* results in significantly more cytosolic Ca^2+^ in the light organ appendages at 24h. Representative images of (A) 24h apo, (B) 18h sym, and (C) 24h sym appendages stained for cytosolic Ca^2+^ with Rhod-3. Scale bar =100 µm. (D) Box plot of the number of puncta present in apo and sym appendages at 18h and 24h when stained with Rhod-3. Letters denote significantly different groups with a p-value < 0.05 from a two-way ANOVA and Tukey post hoc test. Points represent individual appendage puncta counts.

For cytosolic Ca^2+^ to initiate intrinsic apoptosis it has to cause mitochondrial membrane permeabilization. Increased cytosolic Ca^2+^ concentrations lead to mitochondrial Ca^2+^ uptake, which can result in increases in mitochondrial membrane potential via increased NADH production by Ca^2+^-sensitive dehydrogenases (34, 35). However, sustained excess mitochondrial Ca^2+^ concentrations can lead to the loss of mitochondrial membrane potential and the opening of the mitochondrial permeability transition pore (36–38). Therefore, we tested for differences in mitochondrial membrane potential between sym and apo appendages at 24h using the dye tetramethylrhodamine, methyl ester (TMRM). Similar to the sym cytosolic Ca^2+^ staining pattern, we observed significantly more puncta indicative of increased mitochondrial membrane potential in sym appendages relative to apo at 24h (Welch Two Sample t-test, p=0.037) (Figure 8).

**Figure 8.**
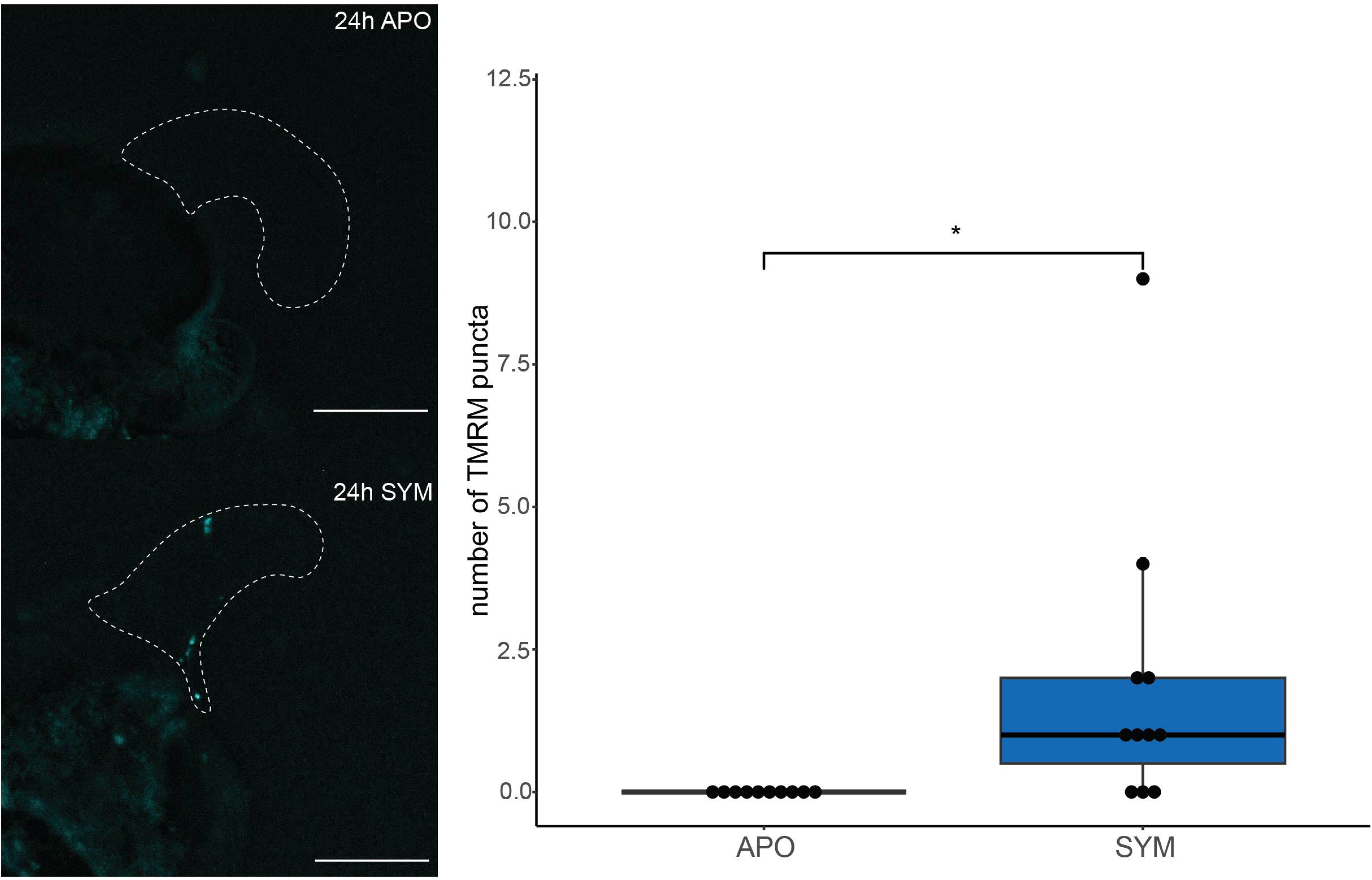
sym appendages have significantly more puncta with high mitochondrial membrane potential relative to apo appendages at 24h. (A) representative image of a 24h apo appendage stained with TMRM (B) representative image of a 24h sym appendage stained with TMRM. Scale bar =100 µm. (C) Box plot of the number of puncta present in apo and sym appendagesg at 18h and 24h when stained with TMRM. * denotes p-value <0.05.

As *aifm1* plays a key role in intrinsic apoptosis and is upregulated in sym appendages vs apo, we used immunocytochemistry to test for nuclear AIF localization indicative of endonuclease activity in sym appendages. Though AIF-immunoreactivity was not observed in nuclei, it was notably more abundant in the CEFs relative to the rest of the light organ, especially in the appendages and along the ciliary ridge (Figure 9). Interestingly, the staining along the ridge is most distinct in apo light organs and becomes more diffuse in sym light organs, particularly at later timepoints (24h, 48h; Figure 9c-f). Negative control light organs in supplemental figure 2 were exposed only to a secondary antibody to ensure fluorescent signal was due to specific binding of the anti-AIF primary antibody.

**Figure 9.**
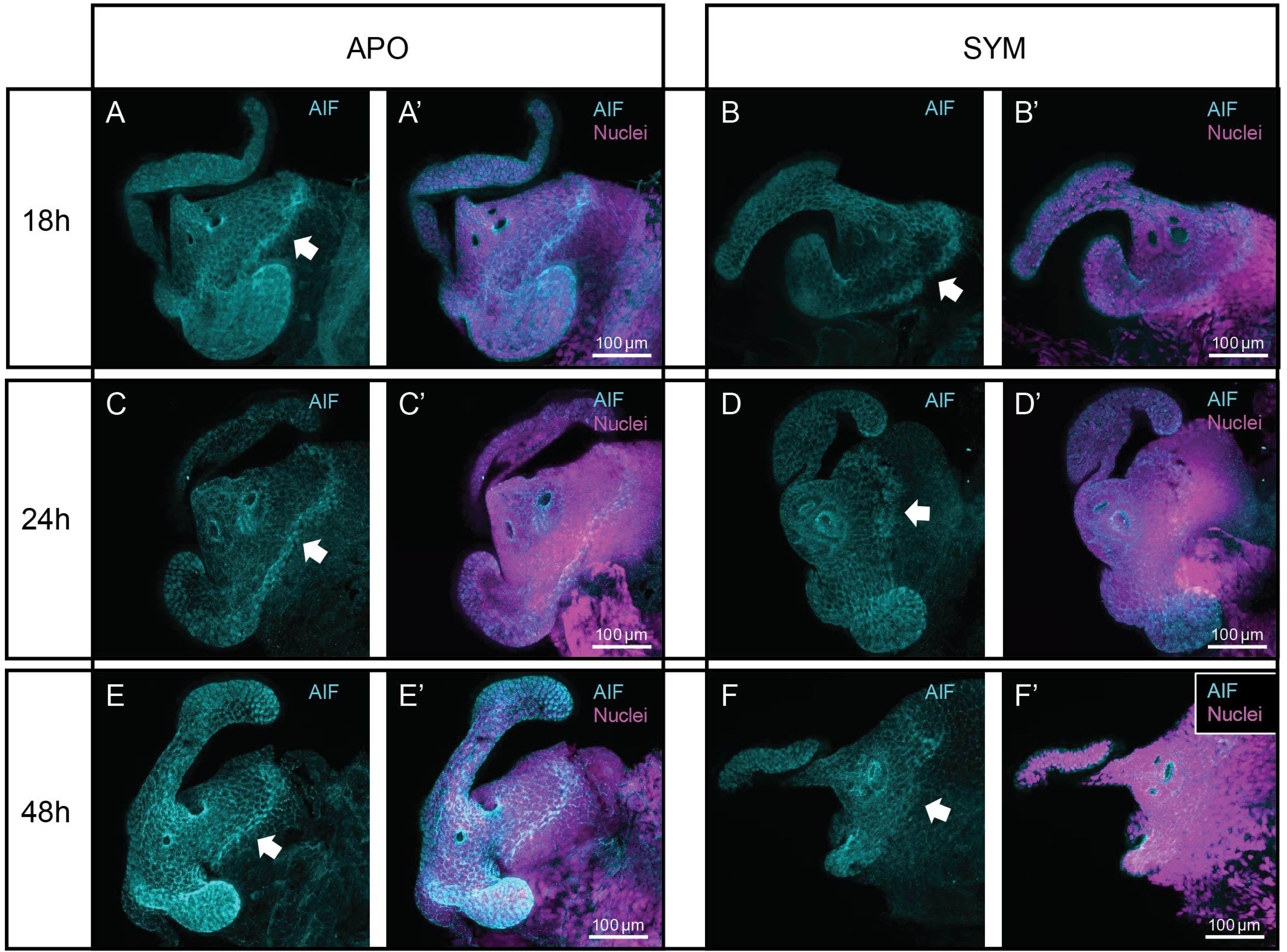
Anti-AIF signal is strongest in the CEF, particularly along the ciliated ridge. Confocal microscopy images of anti-AIF labeling of (A, A’) 18h apo, (B, B’) 18h sym, (C, C’) 24h apo, (D, D’) 24h sym, (E, E’) 48h apo, (F, F’) 48h sym. Anti-AIF is in cyan, nuclei are in magenta, and arrows indicate the ciliated ridge.

## Discussion

Our study builds upon previous work to provide higher resolution into the mechanisms of symbiont-induced developmental apoptosis in the light organ. By isolating appendage gene expression patterns, we gained powerful insights into the cell signaling pathways involved with appendage regression and apoptosis in the CEF. From these data and our subsequent experiments, we propose a model for the apoptotic signaling mediating symbiont-induced developmental apoptosis in the *E. scolopes* light organ, centered around cytosolic Ca^2+^-mediated intrinsic apoptosis (Figure 10). Our findings also provide valuable insight into potential mechanisms of specificity of this apoptosis to the CEF and can be used to narrow down potential signals responsible for initiating this process.

**Figure 10.**
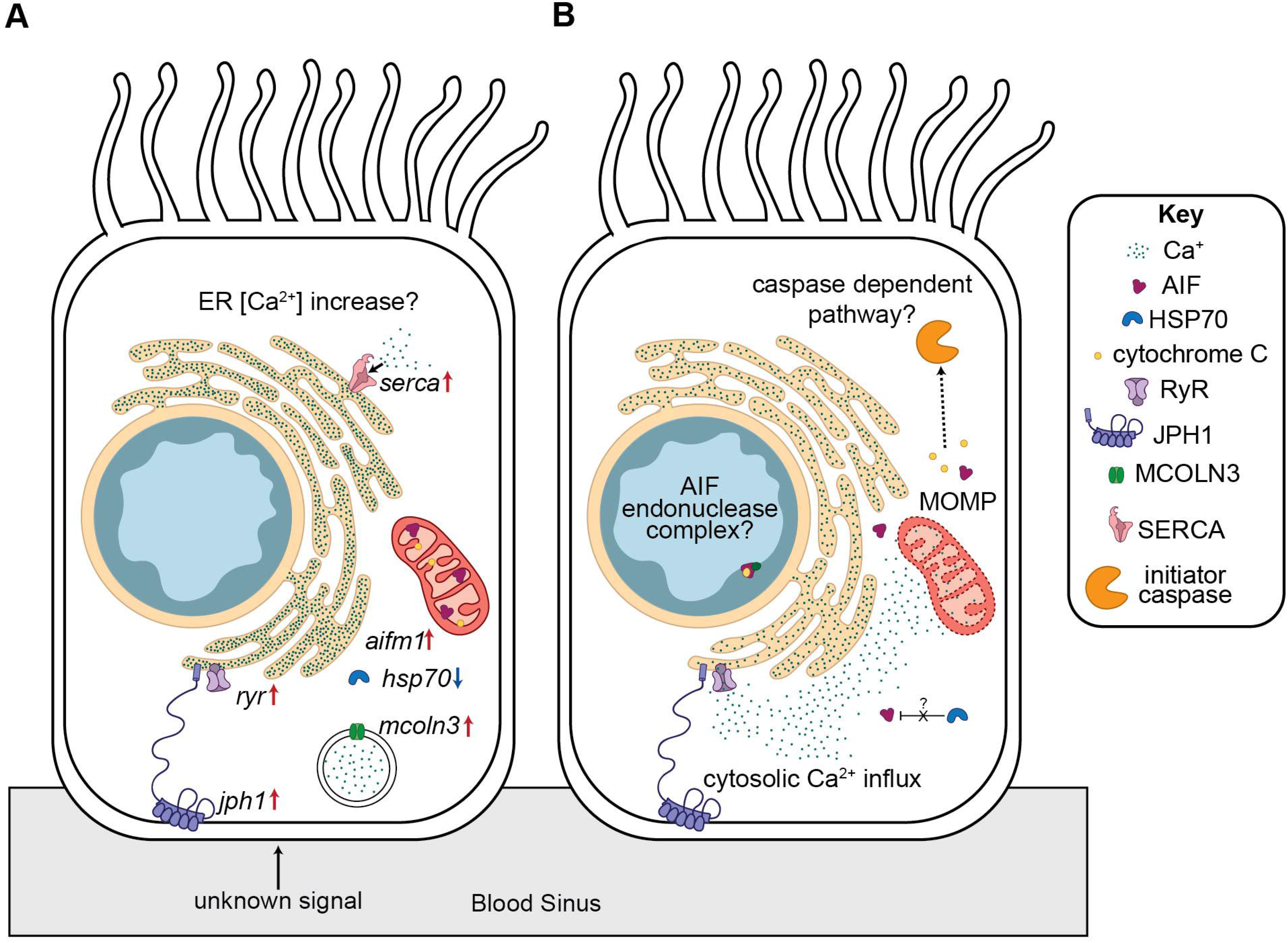
Symbiont induced apoptosis in *E. scolopes* likely occurs via influx of cytosolic Ca^2+^ initiating intrinsic apoptotic signaling. (A) conceptual model of ciliated epithelial cell signaling occurring at 18h that sensitizes the cells to Ca^2+^ mediated apoptosis. Arrows represent up/downregulation of a transcript of interest. (B) conceptual model of the initiation of intrinsic apoptosis via cytosolic Ca^2+^ in the ciliated epithelium.

### Support for cytosolic calcium initiating intrinsic apoptosis in *E. scolopes* appendage regression

We found multiple lines of evidence indicating that excess cytosolic Ca^2+^ initiates intrinsic apoptosis in the light organ appendages as they undergo symbiont-induced morphogenesis (39). In addition to the pathway being upregulated in our GO enrichment analysis, we also found transcriptional evidence for substantial changes to Ca^2+^ transport and homeostasis in sym appendages relative to hatch and apo appendages. The upregulation of conflicting Ca^2+^ transport pathways at 18h in sym appendages likely underlies why we did not consistently observe puncta with increased cytosolic Ca^2+^ at this timepoint. However, by 24h it appears that these conflicting pathways no longer counteract each other, resulting in elevated levels of cytosolic Ca^2+^.

This release may be mediated in part by ryanodine receptors (RyRs), as the large ion channels responsible for releasing Ca^2+^ from the ER into the cytosol are upregulated in sym appendages (Figure 5a). However, RyRs most often act as signal amplifiers, opening in response to already elevated levels of cytosolic Ca^2+^ (33). If this is the case in the sym appendages, it is unclear what induces the initial elevation of cytosolic Ca^2+^. One possibility is endosomal Ca^2+^ release, as mucolipin-3, a Ca^2+^-permeable cation channel that mediates this process, is upregulated in sym appendages relative to both apo and hatch appendages (Supplemental File 2) (40). Alternatively, junctophilin-1 (*jph1*) may mediate RyR channel opening, as it is upregulated in sym appendages relative to both hatch and apo appendages (Figure 6a, supplemental file 3). Junctophilin family proteins facilitate crosstalk between ion channels on the plasma membrane and ion channels on the ER, including RyRs, via the formation of junctional membrane complexes (41, 42). The construction of more of these functional membrane complexes in the ciliated cells of sym appendages could allow for ion channels on the surface of the cells to initiate the opening of RyRs on the ER, releasing Ca^2+^ into the cytosol (42).

Regardless of what initiates ER Ca^2+^ release, respiring mitochondria will uptake cytosolic Ca^2+^ via the mitochondrial Ca^2+^ uniporter (37, 43). The TMRM-stained puncta we observed in sym appendages likely represent cells where mitochondrial membrane potential is temporarily elevated due to NADH production stimulated by increased mitochondrial Ca^2+^ concentrations (34, 35). This increase of membrane potential likely corresponds with an increase in mitochondrial reactive oxygen species (mROS), which are byproducts of aerobic respiration (44). When coupled with the presence of additional signals, such as mROS, sustained excess mitochondrial Ca^2+^ results in depolarization of the inner membrane and the opening of permeability transition pore (37, 38, 45).

Mitochondrial membrane permeabilization is the central event in intrinsic apoptotic signaling, as it initiates the release of pro-apoptotic factors into the cytosol (24, 36, 46). The transcript for one of these pro-apoptotic factors, *aifm1*, is upregulated in sym appendages relative to sym in our RNASeq dataset. In normal conditions AIF resides in the intermembrane space of mitochondria, but it is not immediately released into the cytosol upon mitochondrial membrane permeabilization. Instead, AIF must be cleaved by mitochondrial Ca^2+^-dependent cysteine proteases before it is released into the cytosol (47–49). Within the cytosol AIF can be inhibited by HSP-70 (50). HSP-70 is likely not abundant in the regressing sym appendages, as its transcript is strongly downregulated relative to apo (log-2-fold change < -7) (Figure 6a, supplemental file 2). In the absence of HSP-70, AIF has been shown in other species to translocate to the nucleus to form complexes with endonuclease activity such as the DNA-degradesome complex and the AIF-macrophage migration inhibitory factor (MIF) complex (51, 52). While it remains to be seen if MIF plays a role in apoptosis in the CEF, MIF is known to regulate other aspects of the *E. scolopes*-*V. fischeri* symbiosis (53). As we did not observe any AIF -immunoreactivity in the nuclei of sym appendages, the role of AIF in *E. scolopes* apoptotic signaling is unclear. The lack of nuclear localization could be due to AIFs propensity to form complexes in the nucleus interfering with the antibody’s ability to bind its antigens (51, 52).

Alternatively, the *E. scolopes* AIF may not participate in endonuclease activity and instead induce caspase-dependent cell death, as has been shown in human AIFM3 (54).

### Insights into apoptotic signal specificity

Our data also provide insights as to how the CEFs specifically are targeted for *V. fischeri-* induced apoptosis, given that nearby cells, especially near the ciliated ridge, are likely also exposed to the apoptotic signal. There are multiple lines of evidence suggesting that the cells of the CEFs are primed specifically for cytosolic Ca^2+^-induced intrinsic apoptosis. First, the increased expression of SERCA pumps in sym appendages relative to apo may actually sensitize the CEFs to Ca^2+^ initiated intrinsic apoptosis by increasing Ca^2+^ concentrations in the ER lumen, as a cell’s sensitivity to apoptotic stressors is determined by the combination of concentration of Ca^2+^ in the ER and how closely the mitochondria interface with the ER (32, 39, 55, 56)

Additionally, the spatial localization of AIF within the light organ could prime the CEFs for intrinsic apoptosis. The cells in the appendages and the ciliated ridge have a much higher concentration of AIF signal relative to the rest of the light organ, likely resulting in stronger pro-apoptotic activity upon permeabilization of the mitochondrial outer membrane (57). This specificity may be furthered by the strong downregulation of *hsp-70* in sym appendages but not in the entire sym light organ, as AIF present the cytosol of non-CEF cells would likely be inhibited by HSP-70 (50). Importantly, there is a strong ring of AIF signal along the ciliated ridge, where cells fated to undergo apoptosis are in direct contact with cells that do not undergo symbiont induced post-embryonic developmental apoptosis (9) (9). These neighboring non-ciliated cells likely are exposed to the signal or signals that initiate apoptosis in the CEF and would require mechanisms to not respond to these signals. The limited presence of AIF in non-ciliated cells may be one mechanism by which this symbiont-initiated developmental signal is transduced with high spatial precision.

Another potential mechanism by which non-ciliated cells in the light organ are able survive exposure to the apoptotic signal is the unfolded protein response (UPR) pathway. This is a protective stress response pathway which greatly reduces protein synthesis and upregulates chaperonins to counteract the accumulation of unfolded or misfolded proteins due to ER stress (58, 59). Our data suggest that the UPR is upregulated in sym light organs relative to apo but downregulated in sym appendages vs apo. Upregulation of the UPR in non-appendage light organ cells may serve to bolster their ability to resist apoptotic cues and tolerate host-derived oxidative stress associated with competent *V. fischeri* selection within the light organ (58, 60, 61).

### Potential signals initiating calcium release

The signal following NO attenuation that initiates apoptosis in the CEFs of colonized light organs remains unknown. However, based on our findings that the cells within the CEFs undergo intrinsic apoptosis in response to excess cytosolic Ca^2+^, we can narrow down the potential apoptotic signal for this process. One possibility is that the signal is co-opted from the nervous system, as we see large-scale upregulation of nervous system processes in sym appendages relative to apo and hatch appendages The CEF is likely capable of responding to a nervous system-related apoptotic signal, as excitatory neurotransmitters elicit increased ciliary beat frequency via Ca^2+^ influx in other systems (62–64). Thus, upregulation of DOPA decarboxylase and GO terms related to glutamatergic synaptic transmission, dopamine signaling, and dopamine secretion of in sym appendages relative to apo may be related to the influx of cytosolic Ca^2+^ observed in sym appendages (supplemental file 3) (62, 63). Therefore, sustained exposure to excitatory neurotransmitters could yield the initiation of intrinsic apoptosis specifically in the ciliated epithelia due to their sensitization to Ca^2+^ mediated intrinsic apoptosis via SERCA pump activity (39, 56, 65). Alternatively, non-canonical Wnt/Ca^2+^ signaling may be involved in apoptotic signal transduction, as non-canonical Wnt signaling along with the precursor to protein Wnt-5A, are upregulated in sym appendages relative to apo (66, 67).

## Conclusion

Here we present strong support for the central role of elevated cytosolic Ca^2+^ in symbiont-induced developmental apoptosis in the appendages and CEF of the *E. scolopes* light organ. In addition to the gene expression signatures we found indicative of this process, we show increases in cytosolic Ca^2+^ in sym appendages relative to apo, as well as increases in mitochondrial membrane potential. Together, these data suggest that Ca^2+^ release from the ER initiates intrinsic apoptosis in the appendages and CEF in response to the colonization of the light organ by *V. fischeri*. As Ca^2+^ influx is a known regulator of ciliary beat frequency (62–64) and Ca^2+^ signaling has been shown to be involved in larval metamorphosis in some marine invertebrates (68–70), *E. scolopes* may co-opt existing pathways and machinery to carry out *V. fischeri* induced apoptosis in the post-embryonic light organ.

## Methods

### General Methods

Wild adult *E. scolopes* were collected from Oahu, Hawaii and maintained in facilities at Southern Illinois University (SIU) or Michigan State University (MSU) as previously described (71). Samples in the hatch treatment group were preserved within 3h of hatching. Apo animals were maintained in filter sterilized Instant Ocean (FSIO) while sym animals were incubated with 5000 colony forming units of *V. fischeri* (strain ES114) per mL of FSIO until their respective time points. To confirm successful colonization in sym samples and lack of colonization in apo samples, the squids’ luminescence was measured using a TD-20/20 luminometer. The anesthetic used in all experiments was 2% ethanol in FSIO.

### RNAseq Sample collection and preparation

All samples for RNAseq were collected from January to May of 2021 at SIU. A total of three replicates per treatment group (hatch, 18h apo, 18h sym) were collected, with 25 individuals per replicate. All animals were anesthetized and placed into RNAlater overnight at 4°C before being transferred to -80°C. Samples were then transported to MSU where the anterior appendages of each light organ were removed using forceps. RNA was extracted using the Qiagen RNeasy micro kit (74004). Then, the MSU genomics core tested the quality of the RNA using the Agilent TapeStation system before performing poly-A enrichment, Illumina library preparation, and 35 bp paired end sequencing on an Illumina NextSeq500.

### Transcriptomic data analysis

The *E. scolopes* reference transcriptome used for all analyses was filtered from a previously published assembly (72). Transdecoder (v. 5.7.1) was used to identify contigs with open reading frames (ORFs) and translate these contigs’ longest ORF into amino acid sequences (73). Then, amino acid sequences with a sequence similarity of 98% or more were collapsed using cd-hit (v. 4.8.1) (74). The resulting contigs were extracted from the original assembly, resulting in a reduction in number of contigs from 134,352 to 42,108. The filtered assembly was then annotated with the Swiss-Prot database using Blastx (v. 2.14.) and eggNOG mapper (v.2.1.12) (75–77).

Raw reads were trimmed using Fastp (v. 0.19.5), then mapped and quantified using Salmon (v. 1.10.3) (78, 79). Raw counts were imported into R studio and normalized for length and GC content using TXimport (v. 1.26.1) (80). Prior to downstream analysis, transcripts with an count average below 10 were excluded. Differential gene expression analysis was done using DESeq2 (v. 1.38.3) (model: ∼treatment) (81). Following regularized-log (rlog) transformation a principal component analysis (PCA) was used to detect outliers using the r package PCAtools (82). To visualize expression patterns across treatments, all differentially expressed genes (DEGs) (padj<0.05) from this model were clustered together by expression similarity and visualized using pheatmap (v. 1.0.12). The appropriate contrasts in DEseq2 were used to make pairwise comparisons between all treatments.

The log-2-fold change from the pairwise comparisons was used to calculate rank-based biological process gene ontology (GO) enrichments using GOMWU (29). The GO assignments from the eggNOG annotation were used for this analysis. To summarize the results of these enrichments the R package GO.db (v. 3.16.0) was used to assign all the parent terms associated with the enriched GO terms (padj<0.01). The number of enriched terms per pairwise comparison falling under a set of parent GO terms (GO:0046903, GO:0023052, GO:0003012, GO:0006950, GO:0008152, GO:0007049, GO:0008219, GO:0002376, GO:0006811, GO:0050877, GO:0032502, GO:0044782) were then identified.

### Reanalysis of Moriano-Gutierrez data

Aposymbiotic and symbiotic raw RNAseq reads of whole light organs collected at 24h post hatch generated by Moriano-Gutierrez et al. 2019 were downloaded from NCBI (BioProject PRJNA473394) using SRAtoolkit (30, 83). These were analyzed separately from the appendage RNAseq reads using the pipeline described above.

### Cytosolic Calcium and mitochondrial membrane potential live staining and image analysis

Cytosolic Ca^2+^ was detected using a Rhod-3 imaging kit (Invitrogen R10145) in apoand sym light organs at 18h and 24h. The following modifications were made to the manufacturer’s protocols to ensure the squid tolerated the staining process: FSIO was used as the physiological salt buffer, the loading buffer contained 1.25 mM probenecid, and the incubation buffer contained 250 µM probenecid. The animals were kept in each buffer for 30 minutes at room temperature. Then, the animals were anesthetized with 2% ethanol in FISO, euthanized, and the light organs were dissected for immediate imaging.

Mitochondrial membrane potential was detected in sym and sym light organs at 24h using 250 nM TMRM (Invitrogen I34361). This concentration of dye was not well tolerated by the squid, so the animals were anesthetized, euthanized, and placed into squid ringers (50 mM MgCl_2_, 10 mM CaCl_2_, 10 mM KCl, 530 mM NaCl, 10 mM HEPES). Then the live tissue was stained with TMRM for 30 minutes at room temperature. The presence of ciliary beating was used to confirm that the tissue was kept alive during this process.

For all live staining samples, a single set of appendages were imaged using a Leica thunder stereoscope. Puncta with a minimum particle size of 0.001 within the anterior appendage were quantified using image J following color thresholding (red 90-255) (84). The tissue immediately surrounding the ducts was excluded from analysis, as it showed strong Rhod-3 signal. A two-way ANOVA and subsequent tukey-HSD was used to test for differences in Rhod-3 puncta number between treatments while a Welch’s two sample t-test was used for the TMRM data.

### Localization of AIF in the light organ

To determine the localization of apoptosis-inducing factor 1 (AIF) in the light organ sym and apo light organs at 18h and 24h were extracted and prepared for immunocytochemistry as previously described (15, 85). Briefly, animals were anesthetized and fixed overnight in 4% paraformaldehyde in marine phosphate-buffered saline (mPBS – 50 mM sodium phosphate buffer with 0.45 M NaCl, pH 7.4) at 4 °C. Then, the fixed squid were washed with mPBS and the LOs were removed and permeabilized overnight using 1% Triton-X in mPBS (mPBST). The LOs were then placed in blocking solution (5% bovine serum albumin, 1% normal goat serum in mPBST) for 24h before being incubated with 15 µg/mL of a polyclonal AIF antibody (Invitrogen PA5-48108) for 15 days at 4°C. The percent identity of *E. scolopes* AIF and the human AIF epitope was 64.6% per a Clustal Omega alignment (supplemental figure 3) (86). Following washes with mPBST, samples were incubated overnight at 4°C with 4 µg/mL of secondary antibody (Invitrogen A11008) before the nuclei were stained using TOTO-3. All light organs were mounted in VectaShield and imaged on a Leica Stellaris 5 confocal microscope.

## Code and Data Availability

The raw RNAseq reads generated in this study are available on NCBI (BioProject: PRJNA1492129). The publicly available data used in this study include the *E. scolopes* transcriptome (https://metazoa.csb.univie.ac.at/data/v1/final_assembly_cdhit100.fasta.gz) (72). All code used is available in a Github repository (https://github.com/MaEmery1/Euprymna_scolopes_appendage_apoptosis).

## Acknowledgements

Animals used in these experiments were collected from Hawai i, an indigenous space without which our research would not be possible. We express our gratitude to the Kānaka Maoli (Native Hawaiians) for their hospitality and systems of knowledge, which have sustained Hawai i and allowed generations of scientists to learn from this āina (land) and its people. All imaging was performed at the Michigan State University Center for Advanced Microscopy Core Facility (RRID:SCR_027702). This work was funded by a grant from the National Institute of General Medical Sciences (R35-GM150478).

