## supplemental figure 1 for "Bacterial-driven development is mediated by Calcium-Dependent Intrinsic Apoptosis in the Squid-Vibrio Symbiosis"

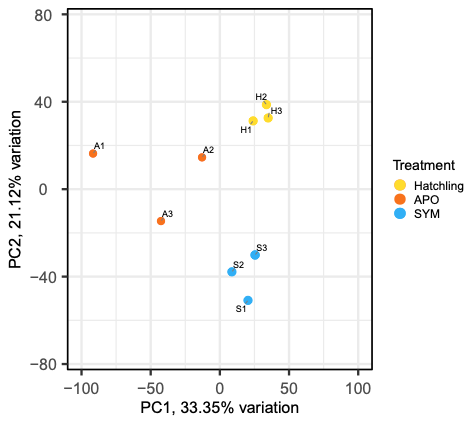

**Supplemental figure 1**

PCA of Rlog normalized expression of all appendage RNAseq samples generated in this study. Abbreviations: APO: aposymbiotic, AYM: symbiotic
