## supplemental figure 2 for "Bacterial-driven development is mediated by Calcium-Dependent Intrinsic Apoptosis in the Squid-Vibrio Symbiosis"

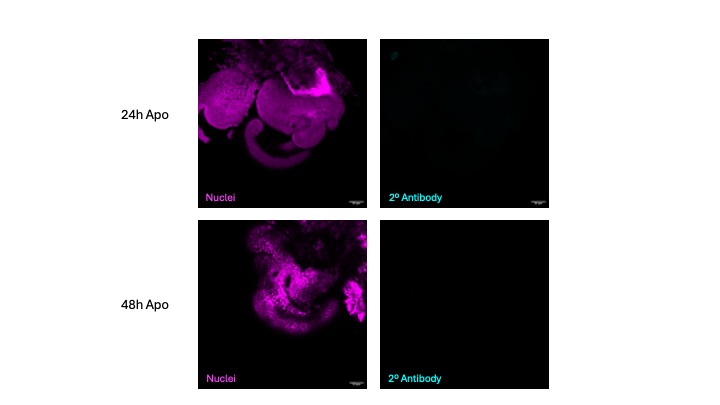

**Supplemental figure 2**

Negative controls for AIF immunocytochemistry, exposed to only secondary antibody. Nuclei are magenta, secondary antibody signal is in cyan.
