## supplemental figure 3 for "Bacterial-driven development is mediated by Calcium-Dependent Intrinsic Apoptosis in the Squid-Vibrio Symbiosis"

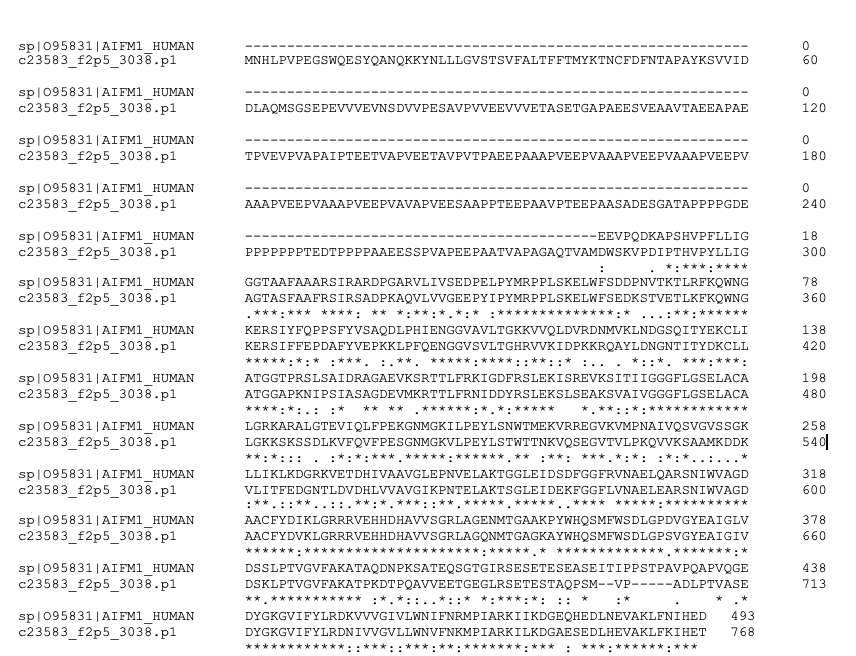

**Supplemental figure 3**

Alignment of the epitope region (121-613) for the polyclonal AIF antibody (Invitrogen PA5-48108) and *E. scolopes* AIF
